# Direct Measurement of Light and Heavy Antibody Chains Using Differential Ion Mobility Spectrometry and Middle-Down Mass Spectrometry

**DOI:** 10.1101/693473

**Authors:** Rafael D Melani, Kristina Srzentić, Vincent R Gerbasi, John P McGee, Romain Huguet, Luca Fornelli, Neil L Kelleher

## Abstract

The analysis of monoclonal antibodies (mAbs) by a middle-down approach is a growing field that attracts the attention of many researchers and biopharma companies. Usually, liquid fractionation techniques are used to separate mAbs polypeptides chains before mass spectrometry (MS) analysis. Gas-phase fractionation techniques such as high-field asymmetric waveform ion mobility spectrometry (FAIMS) can replace liquid-based separations and reduce both analysis time and cost. Here, we present a rapid FAIMS tandem MS method capable of characterizing the polypeptide sequence of mAbs light (Lc) and heavy (Hc) chains in an unprecedented, easy, and fast fashion. This new method uses commercially available instruments and takes ∼ 24 minutes —40-60% faster than regular LC-MS/MS analysis — to acquire fragmentation data using different dissociation methods.

## Main Text

Monoclonal antibodies (mAbs) based on immunoglobulins G (IgGs) are multichain glycoproteins with an approximate molecular weight of 150 kDa. They consist of four polypeptide chains: two light chains (Lc) of ∼ 25 kDa each and two heavy chains (Hc) of ∼ 50 kDa each, all linked together by disulfide bonds^1^. mAbs are the fastest growing class of human therapeutics; the FDA has already approved over 80 therapeutic mAbs since 2018^2^, and they are being used in the treatment of diseases related to cardiovascular, respiratory, hematology, kidney, immunology, and oncology systems^3;4^. As with any other therapeutic, mAbs are required to have their structure well-characterized to ensure drug safety, efficiency, batch-to-batch consistency, and stability over time^5^.

Bottom-up, middle-down, and top-down mass spectrometry (MS) strategies are often used to fulfill and streamline molecular characterization requirements^1^. Bottom-up approaches digest the mAb into peptides before analysis^6^, top-down MS methods analyze intact molecules^7-10^, and middle-down methods are performed by measuring the mass and subsequent fragmenting large pieces or subunits from mAbs (typically 25-50 kDa) that are more suitable for state-of-the-art liquid chromatography tandem mass spectrometry (LC– MS/MS) methods and techniques^11-16^. The subunits or parts of the polypeptide chain can be obtained through the chemical reduction of disulfide bonds—yielding free Lc and Hc^17^— and/or by using a specific enzymatic proteolysis (i.e. digestion with IdeS or IdeZ) that usually generates F(ab’)2 (∼ 100 kDa) and Fc (∼ 50 kDa) pieces^18^ The S-S bounds in these pieces can be further reduced, resulting in three ∼ 25 kDa subunits: one Lc and two portions of Hc named Fc/2 and Fd^19^.

Even the simplest mixture of two unique polypeptide chains obtainable via disulfide bond reduction (i.e., without proteolysis), Lc and Hc, cannot be analyzed effectively by MS without some type of front-end fractionation that can isolate or partially separate the chains. Both the overlap of their charge state envelopes and ionization suppression effects can lower their spectral signal-to-noise ratio (S/N)—particularly for the larger Hc—and could result in co-isolation during fragmentation experiments (tandem MS, or MS/MS). Fractionation methods are typically based on liquid chromatography performed using reverse phase (RP)^17^, size exclusion (SEC)^20; 21^, or ion exchange (IEX)^22; 23^ columns. Each LC-MS/MS run takes several minutes, generally only one fragmentation method is used per run, and multiple injections are needed to maximize sequence coverage^14^. Furthermore, liquid chromatography instruments add expense, with elevated operational costs depending on columns and extent of method development required. Front-end separation based on liquid chromatography also raise issues of sample carryover, contamination, and potential sample losses via irreversible adsorption.

In sharp contrast to liquid-phase separation, a new high-field asymmetric waveform ion mobility spectrometry (FAIMS) device with cylindrical electrodes and improved transmission has recently been described and allows for rapid and effective gas-phase separation of molecules after they are ionized and prior to the mass spectrometer entrance ^24;25^ FAIMS devices operate at atmospheric pressure, conducting ions among an inner and an outer electrode under a high or low electric field^26^. The electric fields are generated from an asymmetric waveform, and the ion separation is based on ion differential mobility. Ions with moderate to no difference in mobility between the high and low fields are conducted to the MS, while ions with a large mobility difference are deflected to the electrodes and are lost. Adding a direct current (DC) voltage—termed the compensation voltage (CV) —to the system alters ion trajectories, which provides a compensation for the drift of specific ions and permits those ions to pass through the electrodes and be analyzed^27; 28^. Ions above ∼ 30 kDa show a strong increase of mobility at high fields, which agrees with expected ion dipole alignment and expands the useful FAIMS separation power^29-31^. Summarizing, changes in CV will favor different groups of ions and function as a filter, as observed for peptides^32^ and proteins^29^. The use of gas-phase fractionation can exclude the liquid separation step for middle-down mAb analysis, making it fast, less expensive, and more robust.

Herein, we present a novel method for fast middle-down analysis of reduced Lc and Hc chains of mAbs without liquid phase pre-fractionation, using only FAIMS Pro™ coupled to an Orbitrap Eclipse™ Tribrid™ mass spectrometer (FAIMS-MS/MS) capable of performing multiple ion fragmentation techniques.

NIST Monoclonal Antibody Reference Material 8671 (NIST), 300 µg, was denatured in 6 M guanidium chloride and reduced using 30 mM tris(2-carboxyethyl)phosphine hydrochloride (TCEP-HCl) for 90 minutes at 37 °C under agitation. The sample was desalted using 3 kDa molecular weight cut-off Amicon Ultra-0.5 (Millipore Sigma), and the solution was buffer exchanged to Optima LC/MS grade water (Fisher Scientific) for over 10 cycles in a refrigerated centrifuge at 4 °C applying 8,000 × *g*. Reduced polypeptide chains were resuspended in 50% acetonitrile containing 0.4% of formic acid for an ∼ 2 µM final concentration.

The polypeptide mixture was sprayed using a Nanospray Flex™ static source (Thermo Fisher Scientific) and medium-length borosilicate-coated emitters (Thermo Fisher Scientific) on an Orbitrap Eclipse™ Tribrid™ (Thermo Fisher Scientific) equipped with FAIMS Pro™ (Thermo Fisher Scientific). The spray voltage was set between 1.5-2.5 kV, and for MS experiments the acquisition range was set between *m/z* 800-2,000 using a resolving power of 7,500 (at *m/z* 200), 2 microscans/spectrum, an average of 20 spectra, 100 ms of maximum injection time, automatic gain control (AGC) target of 5×10^5^ charges, and source collision-induced dissociation (CID) of 10 V; the instrument was operated in “intact protein” mode (pressure of 2 mTorr). FAIMS Pro™ was run at a N_2_ carrier gas flow of 0 L/min, an inner electrode temperature of 100 °C, an outer electrode temperature of 100 °C, a dispersion voltage (DV) of −5,000 V for the asymmetric waveform, an entrance plate voltage of 250 V, and the CV was ranging from −30 to +40 V in 10 V steps.

MS/MS experiments for Lc were carried out at FAIMS CV −20 V using the following parameters: 2 microscans/spectrum, resolving power 120,000 (at *m/z* 200), source CID of 10 V, isolation window of 20 Th centered at *m/z* 1,102 (charge state +21), maximum injection time of 100 ms, AGC target of 5×10^6^ charges, acquisition range set between *m/z* 500-2,000, average of 20 spectra, and ion transfer tube temperature set at 300 °C. For higher-energy collisional dissociation (HCD) normalized collision energy (NCE) was set at 10% for charge state 1, CID NCE at 25% for charge state 1, ultraviolet photodissociation (UVPD) was performed using a 213 nm laser and irradiation time of 70 ms, electron transfer dissociation (ETD) AGC target value for fluoranthene radical anions was set to 7–8×10^5^ charges, default charge state of 3, and ETD reaction times of 5 and 7 ms.

MS/MS experiments for Hc were carried out at FAIMS CV +40 V using the following parameters: 1 microscan/spectrum, resolving power 60,000 of (at *m/z* 200), source CID of 20 V, isolation window of 100 Th centered at *m/z* 1,000 or 1,200 (charge states +49-53 or +41-44 respectively), maximum injection time of 100 ms, AGC target of 5×10^6^ charges, acquisition range set between *m/z* 500-2,000, average of 20 spectra, ion transfer tube temperature set at 300 °C; for HCD, NCE was set at 15% for charge state 1; CID was performed using 10% of NCE for charge state 1; ETD AGC target value for fluoranthene radical anions was set to 7–8×10^5^ charges, default charge state of 3, using ETD reaction times of 2, 5, and 10 ms; electron-transfer/higher-energy collision dissociation (EThcD) was performed using the same ETD conditions with 2 ms reaction time and 15% of NCE for HCD at charge state 1.

The data were analyzed using Thermo XCalibur Qual Browser v4.0.27.10 (Thermo Fisher Scientific) to average spectra and manipulate .raw files. Mass deconvolution of low-resolution data was performed on UniDec GUI v3.0.0^33^. Fragmentation peak fitting and annotation was performed with TDValidator v1.0^14^ (Proteinaceous) using the following parameters: signal to noise (S/N) cutoff of 20 for Lc and 2 for Hc data, max ppm tolerance 20 ppm, sub ppm tolerance 15 ppm, cluster tolerance 0.35, minimum score of 0.7, charge range 1-15, and distribution generator Mercury7. S/N was calculated according to the expression: S/N = (S – B) / (N – B) where S is the signal intensity, B is the spectrum base line intensity, and N is the spectrum noise intensity.

A mixture of reduced Lc and Hc, from NIST mAb reference material, was directly sprayed into an Orbitrap Eclipse™ Tribrid™ mass spectrometer, and the obtained MS spectrum is dominated by the charge state envelope of the Lc with the Hc charge state distribution below 20% of relative intensity (Figure 1A). The constitutional ratio between Lc and Hc for NIST mAb is 1:1. However, S/N for Lc was 119 (*m/z* 1,052, charge state +22), and for Hc it was 14.4 (*m/z* 1,043, charge state +49) and signal intensities were not equivalent. The lower signal and S/N observed for Hc is due to the signal splitting into more charge states than Lc, the presence of more proteoforms (glycosylation), and differences in ionization efficiency. The deconvoluted spectrum (Figure 1B) confirms the abundance discrepancy with Lc representing ∼ 90% of peak intensities while Hc only ∼ 10%. Looking at Hc proteoforms, G1F was the most intense glycoform observed, G0F represented one third of its intensity, and no other Hc glycoform masses were detected.

**Figure 1.**
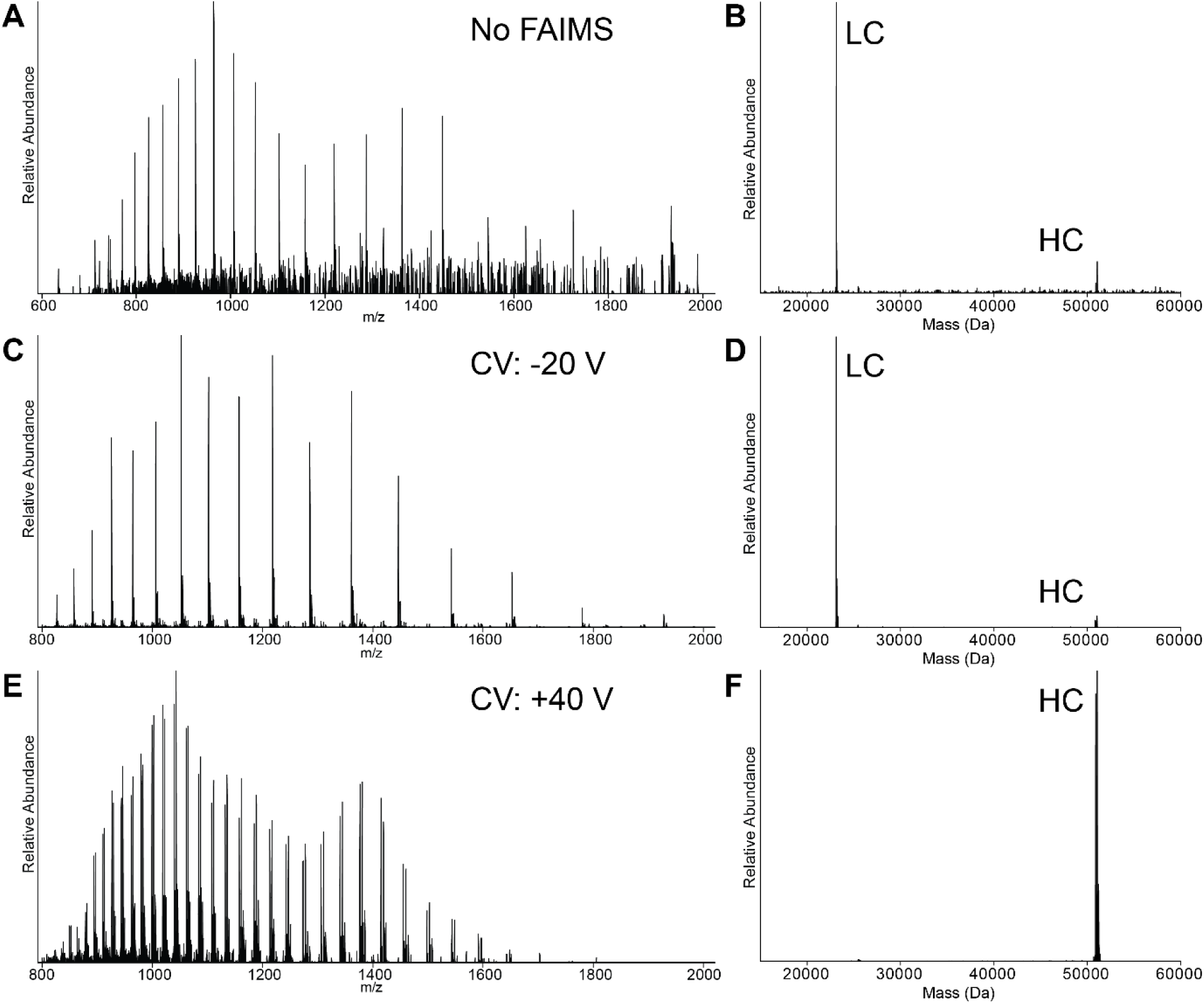
Mass spectra of reduced mAb (obtained from NIST) for two different settings of the FAIMS compensation voltage (CV) compared to no FAIMS. Spectra from direct injection (no FAIMS) of the mixture containing reduced light (Lc) and heavy (Hc) chains with the resultant spectra displayed in the *m/z* domain (A) and the deconvoluted spectra in the mass domain (B). Spectra obtained using FAIMS and applying −20 V of CV and displayed in the *m/z* domain (C) or deconvoluted into the mass domain (D) spectra. Spectra obtained using FAIMS and applying +40 V of CV and displayed in the *m/z* domain (E), or deconvoluted into the mass domain (F).

Spraying the same mixture into the instrument equipped with FAIMS Pro™ and stepping the compensation voltage (CV) by 10 V from −30 V to +40 V allowed the gas-phase separation of Lc and Hc based on their ion mobilities across the generated high field asymmetric waveforms (Supplementary Figures S1 and S2). Acquisition was performed for three minutes in each CV 10 V steps, and the −20 V step presented the clearest spectrum for Lc (Figure 1C). The most abundant charge state showed a S/N of 2,390 for *m/z* 1,052 (charge state +22), which is equal to an increase of 20-fold compared to the no FAIMS spectrum. Moreover, the Hc charge distribution observed was below 3% of total ion relative intensity. The deconvoluted spectra is composed of 95.2% of Lc and 4.8% of Hc based on peak intensities (Figure 1D), and the Lc observed average mass of 23,127.31 Da is −11.2 ppm off the theoretical mass 23,127.57 Da. Adducts of sodium (22 Da), guanidine (60 Da), their combination, and the double guanidine species were also observed as well as a 162 Da mass shift which corresponds to the addition of one hexose (Hex) to Lc (Figure 2A). The non-enzymatic, but covalent adduction of a Hex sugar molecule on a lysine or on a protein N-terminus is called “glycation”, and for the Lc of NIST mAb there are 22 distinct glycated peptides reported^34^. Lc+Hex is reported as trace level post-translational modifications (PTMs)^35^. In addition, the glycated Lc proteoforms correspond to ∼ 4% of the total ion intensity of the non-modified Lc, indicating that FAIMS-MS is suitable to detect low stoichiometry mAbs PTMs.

**Figure 2.**
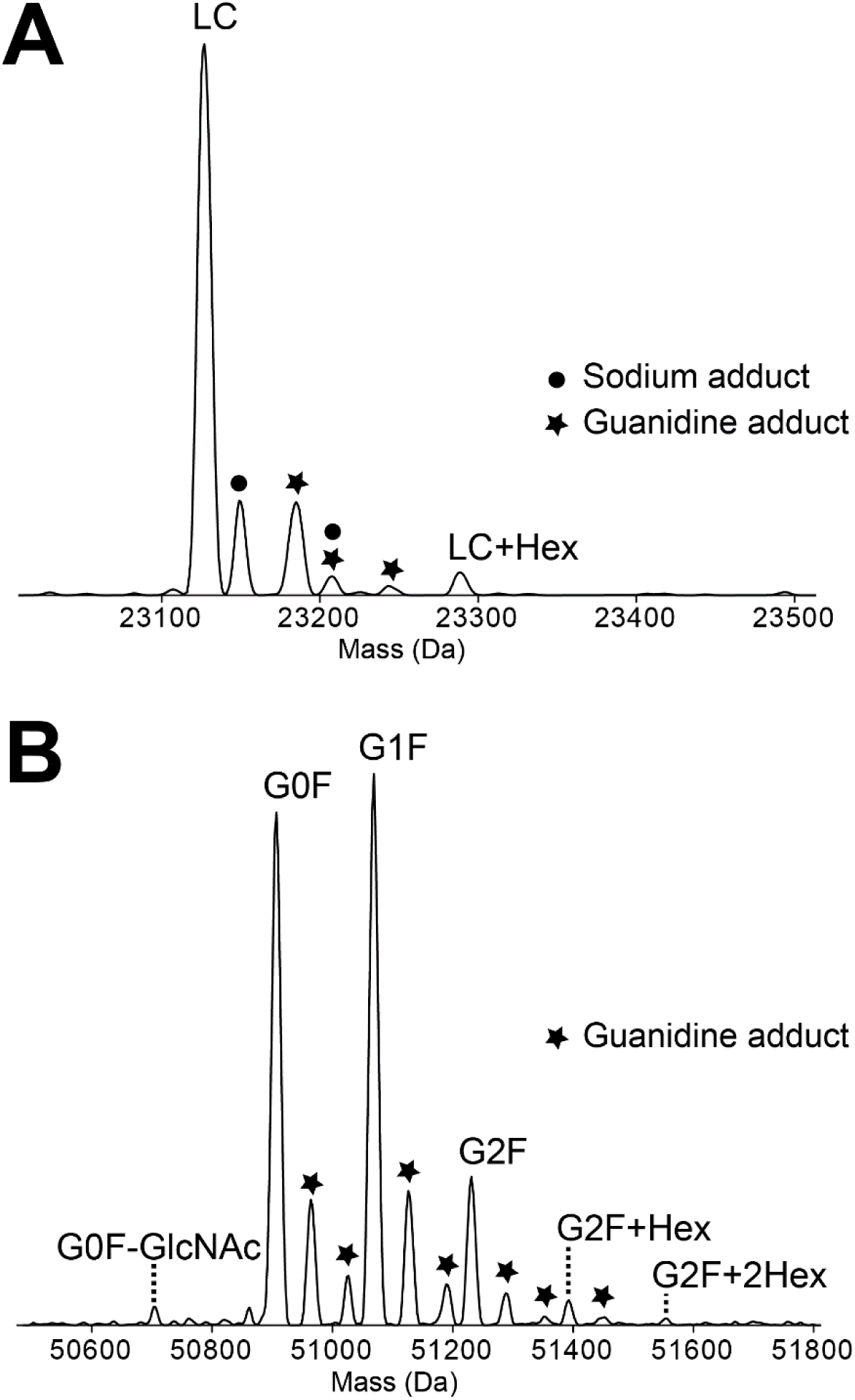
Expanded section of deconvoluted spectrum of light (Lc) and heavy (Hc) chains. Spectra obtained applying −20 V and +40 V of CV were deconvoluted and zoomed-in to show Lc (A) and Hc (B) proteoforms. Sodium adducts (+22 Da) are represented by a circle (•) and guanidine adducts (+60 Da) by a star (★). Addiction of hexose (+162 Da) is characterized by +Hex and the loss of *N*-acetylglucosamine (−203 Da) as -GlcNAc.

On the other hand, the CV of +40 V generated the cleanest spectrum for Hc (Figure 1E) with no detection of Lc and a S/N of 477 for *m/z* 1,043 (charge state +49), which corresponds to a 34-fold increase compared to the spectrum recorded without FAIMS. The observed deconvoluted average mass for Hc G1F was 51,068.59 Da, −10.4 ppm off from the theoretical mass 51,069.12 Da. In the deconvoluted spectrum only Hc was detected (Figure 1F), and it presented 6 glycoforms (Figure 2B): G0F-GlcNAc, G0F, G1F, G2F, G2F+Hex, and G2F+2Hex. Their relative abundances based on ion intensity are 1.5%, 40.5%, 43.6%, 11.8%, 2.1%, and 0.5% respectively. Single- and double-guanidine (60 Da) adducts were also observed. The observed ratios of the glycoforms are in accordance with the literature findings^36;37^. As G0F-GIcNAc, G2F+Hex, and G2F+2Hex were just not seen without the use of FAIMS due to their low abundance. Their detection and accurate relative quantitation using FAIMS are a good indicator of the heightened sensitivity afforded using FAIMS-MS.

Preventing the overlap of Lc and Hc charge state envelopes in the *m/z* space permits the isolation of a single charge state of each polypeptide chain or the isolation of multiple charge states of the same polypeptide chain without co-isolation with the other chain. Avoiding co-isolation is important for successful fragmentation of a single species, generating non-chimeric spectra, which are subsequently easier to correctly interpret and match against the polypeptide primary chain sequence. A single charge state of the Lc chain was quadrupole-isolated using a 20 Th window and fragmented by HCD, CID, ETD, and UVPD. MS/MS data were acquired for only three minutes in each dissociation method, and Lc graphical fragmentation maps were generated using TDValidator (Supplementary Figure S3). The combination of all the quickly acquired dissociation maps (resulting from 12 minutes. total instrument time) yielded 65% sequence coverage for Lc (Figure 3A). For Hc, multiple charge states were isolated in a 100 Th isolation window, and ions were fragmented using HCD, CID, ETD, and EThcD (Supplementary Figure S4). Acquisition time was comparable to the one needed for the Lc, and the combination of fragmentation maps resulted in 34.4% of sequence coverage for the most abundant proteoform Hc G1F (Figure 3B). No changes in the fragmentation maps were observed considering G0F or G2F glycans.

**Figure 3.**
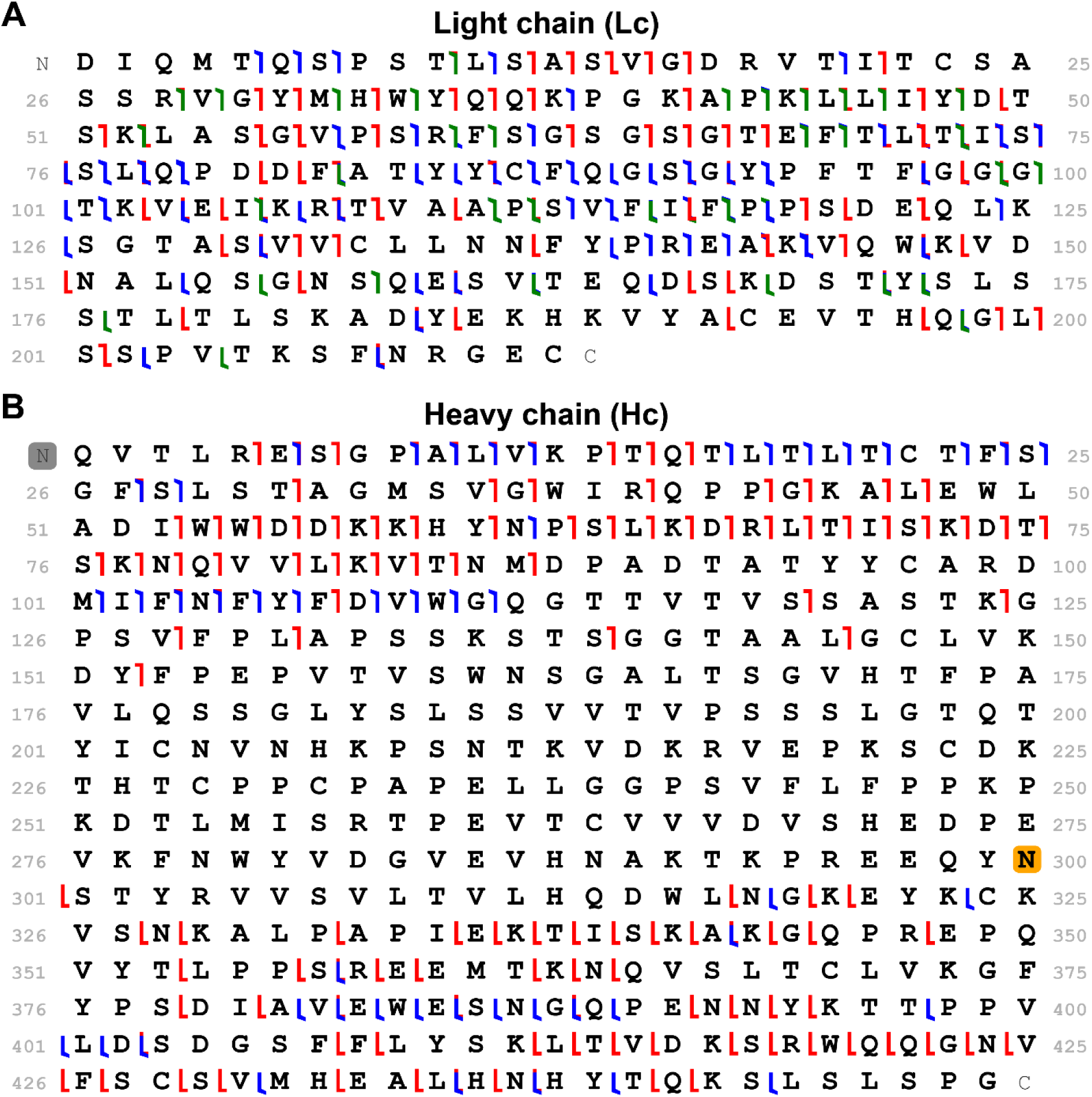
Graphical fragmentation maps obtained from middle-down tandem MS of light (Lc) and heavy (Hc) chains. Cumulative fragmentation maps obtained from HCD, CID, ETD, and UVPD dissociation methods applied on Lc (A) and from HCD, CID, ETD, and EThcD applied to Hc (B). The red brackets represent *c-* and *z-* ions, blue brackets *b-* and *y-* ions, and green brackets *a-* and *x-* ions. The gray rectangle denotes a pyroglutamic acid post-translational modification and the orange rectangle the addition of the *N*-linked glycan G1F mass (the most abundant proteoform observed for the standard mAb obtained from NIST).

FAIMS-MS/MS was capable of separation and analysis of a mixture of reduced Lc and Hc from mAb in an unprecedented, easy, and fast fashion without the use of liquid chromatography or any other liquid fractionation technique. All fragmentation data obtained correspond to a minimum of four regular LC-MS/MS runs which would take around 10-15 minutes each, totaling 40-60 minutes of analysis not considering loading times, blanks, washes, and standards. The FAIMS-MS/MS method presented here required only 24 minutes to acquire all fragmentation data on Lc or Hc using 4 different dissociation methods, which is 40-60% faster than regular LC-MS/MS analysis. Further improvements in the acquisition routine can make the method even faster for the characterization of mAb and mAb conjugates. In this work, samples were manually sprayed, and data was acquired directly from Tune (instrument controller software). However, it is possible to use automated nanospray or microspray to run FAIMS-MS/MS in a high throughput manner far simpler than current LC-MS/MS analyses. The analysis of mAbs is a growing field that attracts the attention of many researchers and biopharmaceutical companies, and the new FAIMS-MS/MS method presented can improve speed, limit artefacts, and reduce costs of middle-down mAb analysis.

## Abbreviations

mAb: Monoclonal Antibody
IgGs: Immunoglobulins G
Lc: Light Chain
Hc: Heavy Chain
FDA: Food and Drug Administration
MS: Mass Spectrometry
LC–MS/MS: Liquid Chromatography Tandem Mass Spectrometry
S/N: Signal to Noise
MS/MS: Tandem Mass Spectrometry
RP: Reverse Phase
SEC: Size Exclusion
IEX: Ion Exchange
FAIMS: High-field asymmetric waveform ion mobility spectrometry
DC: Direct Current
CV: Compensation Voltage
TECP-HCl: (2-carboxyethyl)phosphine Hydrochloride
AGC: Automatic Gain Control
CID: Collision Induced Dissociation
DV: Dispersion Voltage
HCD: Higher-energy Collisional Dissociation
NCE: Normalized Collision Energy
UVPD: Ultraviolet Photodissociation
ETD: Electron Transfer Dissociation
EThc: DElectron-transfer/Higher-energy Collision Dissociation
Hex: Hexose
PTMs: Post-translational Modifications

## Acknowledgments

This research was carried out in collaboration with the National Resource for Translational and Developmental Proteomics under Grant P41 GM108569 from the National Institute of General Medical Sciences (NLK) and supported by the Sherman Fairchild Foundation. LF would like to thank the University of Oklahoma, Department of Biology, for start-up funds

## Disclosure of potential conflicts of interest

KS and RH are Thermo Fisher Scientific employees, and NLK declares a COI with Proteinaceous.

