## Supplemental File for "Direct Measurement of Light and Heavy Antibody Chains Using Differential Ion Mobility Spectrometry and Middle-Down Mass Spectrometry"

### Supplementary Figures

**Supplementary Figure S1.** Mass spectra of reduced NIST mAb direct injection applying different settings of FAIMS compensation voltage (CV).

**Supplementary Figure S2**. Deconvoluted mass spectra of reduced NIST mAb direct injection applying different settings of FAIMS compensation voltage (CV).

**Supplementary Figure S3.** Graphical fragmentation maps of different dissociation methods obtained from middle-down tandem MS of NIST mAb light chain (Lc).

**Supplementary Figure S4.** Graphical fragmentation maps of different dissociation methods obtained from middle-down tandem MS of NIST mAb heavy chain (Hc).

**Supplementary Figure S1.** Spectra obtained from direct injection of the mixture containing reduced light (Lc) and heavy (Hc) chains from NIST mAb stepping FAIMS compensation voltage (CV) by -10 V from +40 V to -30 V. Spectra are displayed in the *m/z* domain and CV +40 V and -20 V presented the clearest spectrum for Hc and Lc, respectively.

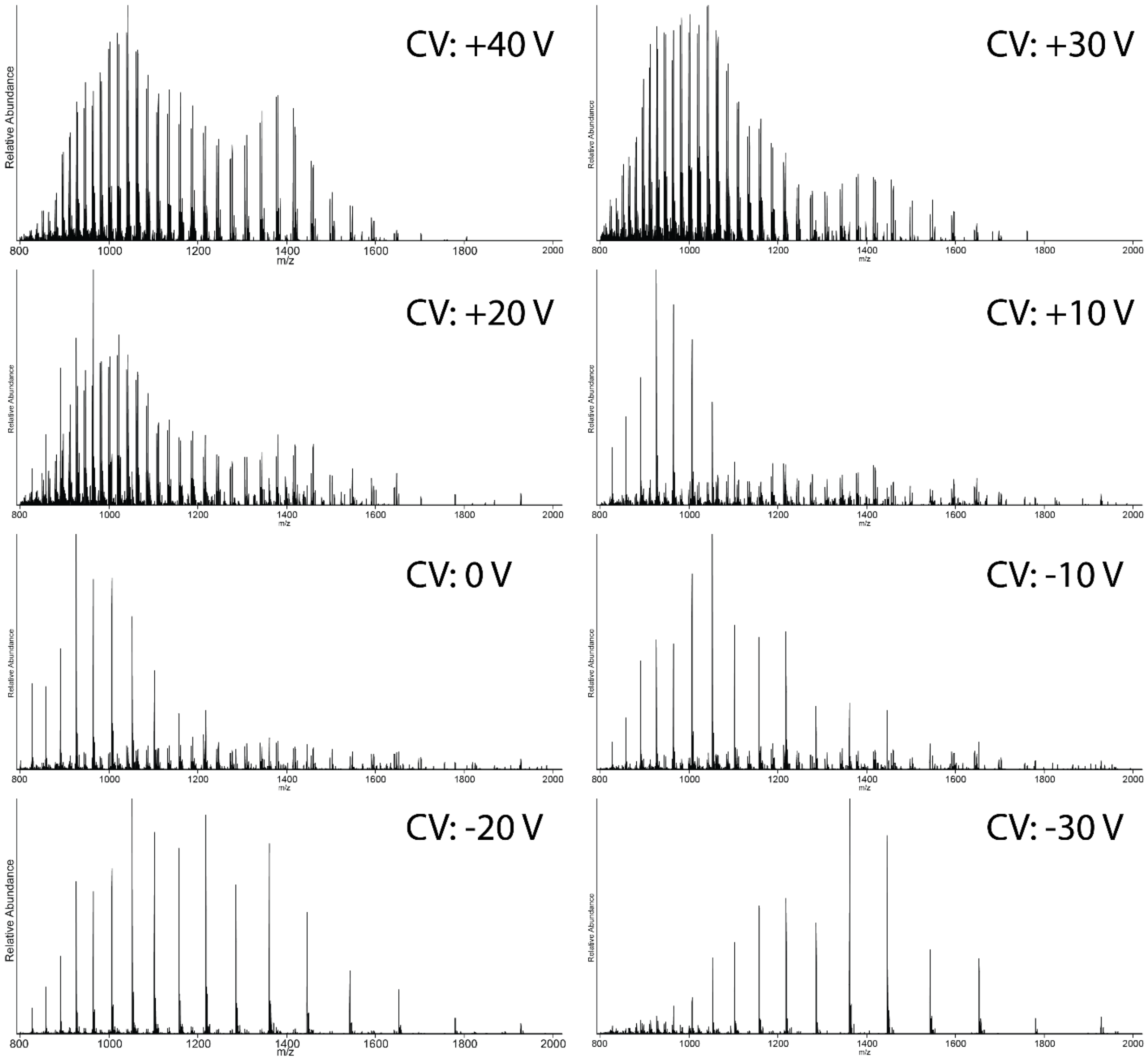

**Supplementary Figure S2.** Deconvoluted spectra obtained from direct injection of the mixture containing reduced light (Lc) and heavy (Hc) chains from NIST mAb stepping FAIMS compensation voltage (CV) by -10 V from +40 V to -30 V. Spectra are displayed in mass domain and CV +40 V and -20 V presented the clearest spectrum for Hc and Lc, respectively.

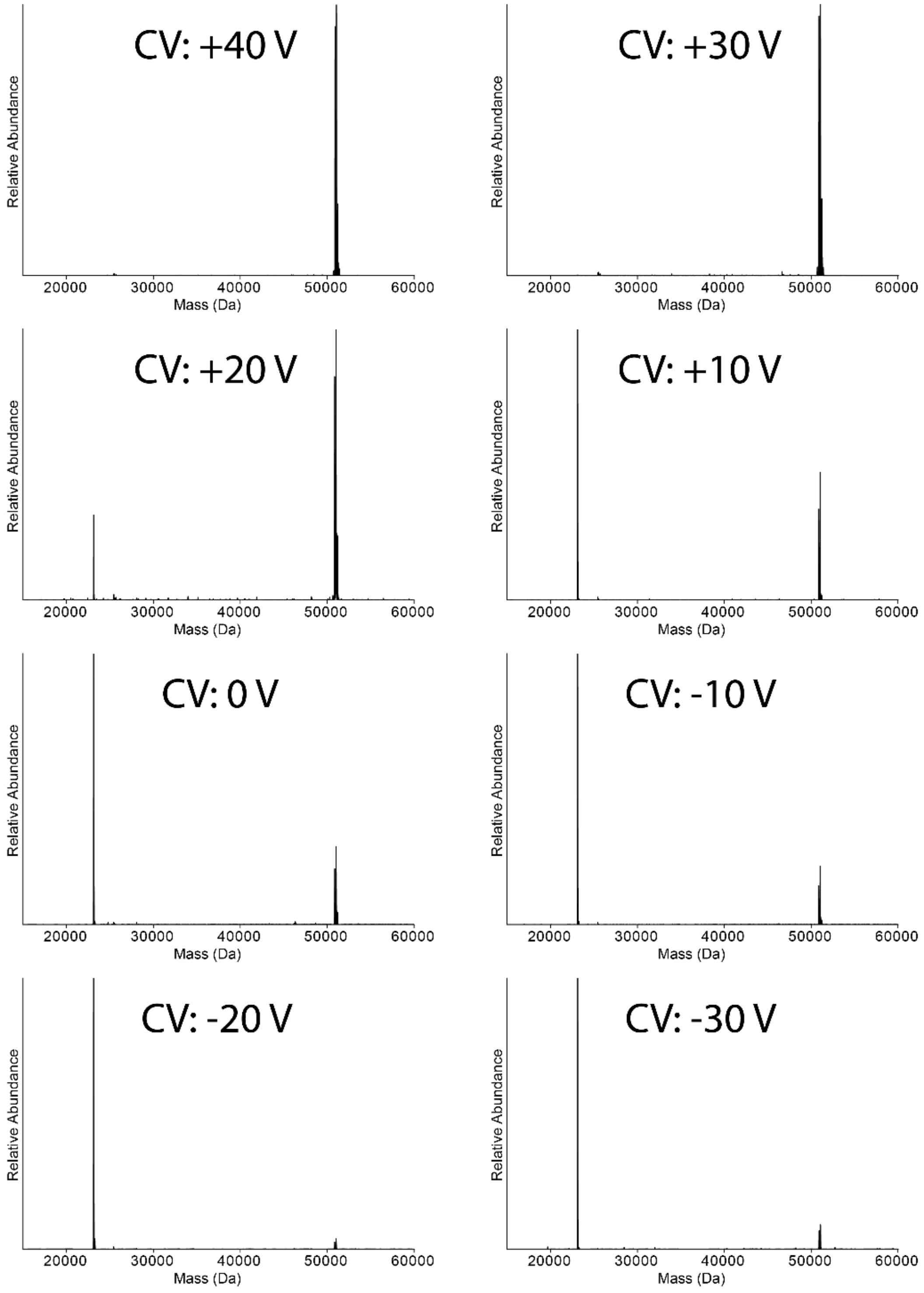

**Supplementary Figure S3.** Graphical fragmentation maps for NIST mAb light chain (Lc) obtained using (A) HCD, (B) CID, (C) ETD, and (D) UVPD dissociation methods. The red brackets represent *c*- and *z*- ions, blue brackets *b*- and *y*- ions, and green brackets *a*- and *x*- ions.

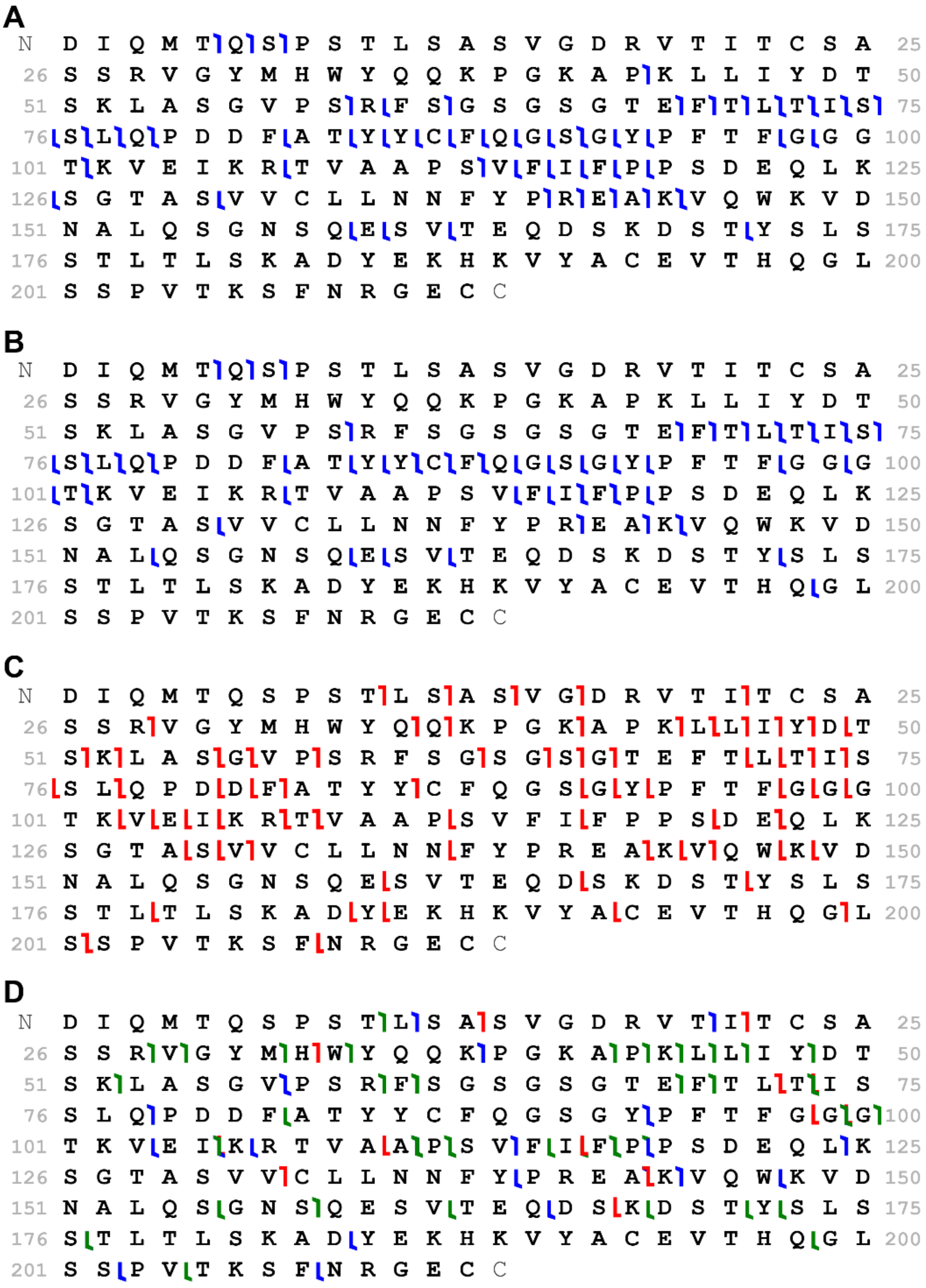

**Supplementary Figure S4.** Graphical fragmentation maps for NIST mAb heavy chain (Hc) obtained using (A) HCD, (B) CID, (C) ETD, and (D) EThcD dissociation methods. The red brackets represent *c*- and *z*- ions, blue brackets *b*- and *y*- ions, and green brackets *a*- and *x*- ions. The gray rectangle denotes a pyroglutamic acid post-translational modification and the orange rectangle the addition of the N-linked glycan G1F mass.

A

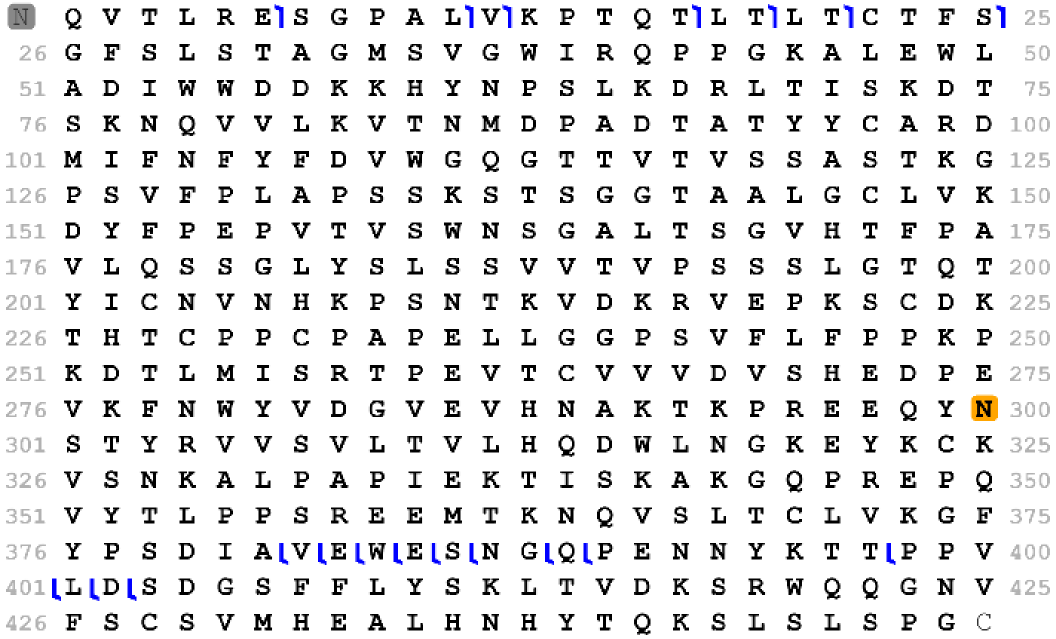


B

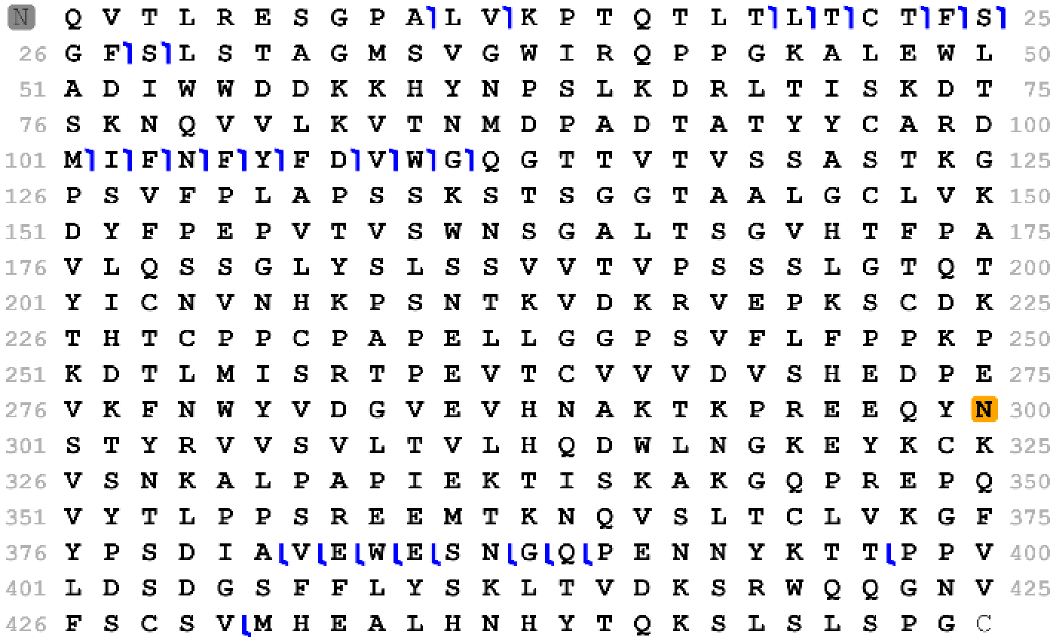

**C**

N Q V T L R E S G P A L V K P T Q T L T L T C T F S 25
26 G F S L S T A G M S V G W I R Q P P G K A L E W L 50
51 A D I W W D D K K H Y N P S L K D R L T I S K D T 75
76 S K N Q V V L K V T N M D P A D T A T Y Y C A R D 100
101 M I F N F Y F D V W G Q G T T V T V S S A S T K G 125
126 P S V F P L A P S S K S T S G G T A A L G C L V K 150
151 D Y F P E P V T V S W N S G A L T S G V H T F P A 175
176 V L Q S S G L Y S L S S V V T V P S S S L G T Q T 200
201 Y I C N V N H K P S N T K V D K R V E P K S C D K 225
226 T H T C P P C P A P E L L G G P S V F L F P P K P 250
251 K D T L M I S R T P E V T C V V V D V S H E D P E 275
276 V K F N W Y V D G V E V H N A K T K P R E E Q Y N 300
301 S T Y R V V S V L T V L H Q D W L N G K E Y K C K 325
326 V S N K A L P A P I E K T I S K A K G Q P R E P Q 350
351 V Y T L P P S R E E M T K N Q V S L T C L V K G F 375
376 Y P S D I A V E W E S N G Q P E N N Y K T T P P V 400
401 L D S D G S F F L Y S K L T V D K S R W Q Q G N V 425
426 F S C S V M H E A L H N H Y T Q K S L S L S P G C

**D**

N Q V T L R E S G P A L V K P T Q T L T L T C T F S 25
26 G F S L S T A G M S V G W I R Q P P G K A L E W L 50
51 A D I W W D D K K H Y N P S L K D R L T I S K D T 75
76 S K N Q V V L K V T N M D P A D T A T Y Y C A R D 100
101 M I F N F Y F D V W G Q G T T V T V S S A S T K G 125
126 P S V F P L A P S S K S T S G G T A A L G C L V K 150
151 D Y F P E P V T V S W N S G A L T S G V H T F P A 175
176 V L Q S S G L Y S L S S V V T V P S S S L G T Q T 200
201 Y I C N V N H K P S N T K V D K R V E P K S C D K 225
226 T H T C P P C P A P E L L G G P S V F L F P P K P 250
251 K D T L M I S R T P E V T C V V V D V S H E D P E 275
276 V K F N W Y V D G V E V H N A K T K P R E E Q Y N 300
301 S T Y R V V S V L T V L H Q D W L N G K E Y K C K 325
326 V S N K A L P A P I E K T I S K A K G Q P R E P Q 350
351 V Y T L P P S R E E M T K N Q V S L T C L V K G F 375
376 Y P S D I A V E W E S N G Q P E N N Y K T T P P V 400
401 L D S D G S F F L Y S K L T V D K S R W Q Q G N V 425
426 F S C S V M H E A L H N H Y T Q K S L S L S P G C
